# External drivers of BOLD signal’s non-stationarity

**DOI:** 10.1101/2021.09.07.459325

**Authors:** Arian Ashourvan, Sérgio Pequito, Maxwell Bertolero, Jason Z. Kim, Danielle S. Bassett, Brian Litt

## Abstract

A fundamental challenge in neuroscience is to uncover the principles governing how the brain interacts with the external environment. However, assumptions about external stimuli fundamentally constrain current computational models. We show in silico that unknown external stimulation can produce error in the estimated linear time-invariant dynamical system. To address these limitations, we propose an approach to retrieve the external (unknown) input parameters and demonstrate that the estimated system parameters during external input quiescence uncover spatiotemporal profiles of external inputs over external stimulation periods more accurately. Finally, we unveil the expected (and unexpected) sensory and task-related extra-cortical input profiles using functional magnetic resonance imaging data acquired from 96 subjects (Human Connectome Project) during the resting-state and task scans. Together, we provide evidence that this embodied brain activity model offers information about the structure and dimensionality of the BOLD signal’s external drivers and shines light on likely external sources contributing to the BOLD signal’s non-stationarity.

## 1 Introduction

Over the past few decades, functional MRI has widened our understanding of the functional organization of intrinsic brain networks and their role in cognition and behavior. Classical univariate (i.e., voxel-wise) analyses of fMRI signal (i.e., blood-oxygenation level-dependent, or BOLD) have been instrumental in probing the specialized function of brain regions. More recent approaches using functional connectivity and network neuroscience portray a complex and multi-scale set of interactions between brain structures. Following this view, a wide array of graph theoretical and complex systems tools have been used to describe BOLD dynamics^1;2;3^.

Despite these efforts, we still lack a unified mechanistic framework that overcomes three key limitations. First, the features of the BOLD signal that are important for neural activity are unclear. Several prior studies demonstrate a relation between BOLD and slow amplitude features of cortical activity^4;5;6^, and between BOLD and the hemodynamic response function (HRF)^7;8^. These studies imply that the low frequency component of the BOLD signal contains information relevant to underlying neural dynamics^9;10^, although it is also clear that the signal contains artifact^11;12^. Due to the mixture of signal and artifact in the BOLD time series, it is possible that the common practice of band-pass filtering the BOLD signal at low frequencies may exclude functionally relevant signal^13;14^. Second, many graph theoretic and network analyses are inherently descriptive in nature, and lack the power to give a generative understanding of the relationship between model inputs and outputs (for extensions of these approaches that move beyond description into explanation and prediction, see^15^). Finally, model-based approaches often treat the brain as an isolated system by ignoring external input, or assuming an artificial profile of internal and external noise.

To address these three limitations, we develop a generative framework that explicitly includes exogenous input (e.g., external sensory or subcortical structures’ inputs), and provide evidence that the brain’s activity can be fruitfully understood in the context of its natural drivers. Specifically, we use a multivariate autoregressive model with unknown inputs to capture the spatiotemporal evolution of the BOLD signal driven by extra-cortical inputs. These models have been used to characterize and predict the evolution of several synthetic and biological systems^16;17;18;19^. For instance, Chang and colleagues (2012) leveraged a multivariate linear dynamical system’s framework and the patients’ intracranial EEG to model the cortical impulse response to the direct electrical stimulation. Many prior studies use this^20^ and similar methods such as Granger causality and dynamic causal modeling (DCM) for understanding the directed functional connectivity of BOLD^21;1;22;23^. While some prior studies account for the effect of exogenous input^1;24^, they typically assume a simple known and abstract form of the input function^19^. Moreover, the inability of models such as DCM to capture signal variations beyond those caused by the external inputs makes the connectivity estimation highly dependent on the assumed number and form of the inputs^25^.

In this work, we treat the exogenous inputs to the cortex as *unknown* parameters of a linear time-invariant (LTI) system, which we estimate following recent developments in linear systems theory^26^. We use these developments to provide new insights into how the brain responds to ongoing task requirements, and to shine a light on factors that contribute to the dynamics of cortical functional connectivity. To demonstrate our approach’s utility, we begin with a proof-of-concept where we consider synthetic examples for which we retrieve the external inputs’ spatiotemporal profiles of a known LTI system. We demonstrate that unknown external inputs result in apparent changes in internal system parameters, and consequently, in estimated external inputs’ error. Also, we show that using internal system parameters estimated from time windows without external stimulation significantly improves our ability to extract external inputs’ profile from periods with external stimulation, expect for simulations with relatively low external inputs and signal-to-noise.

Next, we test the hypothesis that variations in cortical dynamics during different tasks or cognitive states can be accurately modeled as external excitations on fairly stable interactions between cortical regions. Specifically, we recover the unknown external cortical inputs during resting-state and task scans for 96 subjects with the lowest motion artifact from the Human Connectome Project (HCP). Our results demonstrate that using system parameters estimated from resting-state scans enables uncovering the expected spatiotemporal profiles of external sensory (i.e., visual cues) and task-related extra-cortical inputs, while system parameters estimated from task scans result in highly inaccurate input estimations. In addition, an in-depth examination of estimated inputs during task scans reveals the spatiotemporal patterns of other task-related inputs that were not captured by the abstract task regressors.

Lastly, we measure the non-stationarity of estimated external inputs over resting-state scans to examine the assumption of the system’s time-invariance and to identify exogenous determinants of the BOLD signal’s non-stationarity. Recently, the nature of non-stationarity of BOLD signal and dynamic functional connectivity has been a topic of scientific debate, as several recent publications paint seemingly contrasting portraits of the processes’ stationarity underlying the brain’s functional dynamics^27;28;29;30;31;32^. However, to the best of authors’ knowledge, no study examines the BOLD signal’s stationarity in the context of time-varying external inputs and their effects. Our results show that the inputs to several brain regions, most notably over default mode network, estimated from the resting-state scans display significantly high non-stationarity compared to other brain regions. Together, we demonstrate that our framework allows us to uncover spatiotemporal patterns and dimensionality of unknown cortical drivers. These findings offer insight into how a relatively static relation between brain regions and exogenous drivers can give rise to complex cortical dynamics and contribute to their non-stationarity.

## 2 Materials and Methods

### 2.1 Linear time-invariant (LTI) dynamical systems with external inputs

Each region *i* of interest (ROI) from which the BOLD signal is collected provided us with a time series described by *x_i_*[*k*] at sampling point *k* = 0,…, *T*. A total of *n* = 100 regions are considered and the collection of these signals is captured by the vector *x*[*k*] = [*x*_1_[*k*] … *x_n_*[*k*]]^T^, with *k* = 0,…, *T*, which we refer to as the *state of the system* (i.e., it describes the evolution of the BOLD signal across different regions). The evolution of the system’s state is mainly driven by (*i*) the cross-dependencies of the signals in different regions (not necessarily adjacent), and (*ii*) the external inputs that are either excitation noise or inputs arriving from the environment surrounding the regions captured by the state of the system (e.g., stimulus arriving from subcortical structures not accounted for during BOLD signal collection).

Subsequently, a first step towards modeling the evolution of the system’s state is:

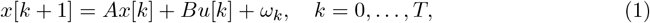

where 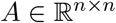 described the autonomous dynamics, 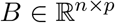 is the input matrix that describes the impact of inputs (i.e., external drivers) 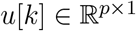 on the system state’s evolution, and 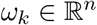 is the internal dynamics noise (i.e., internal drivers) at sampling point *k*. Notice that 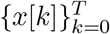 is the BOLD signal at the different ROIs and is the only known. However, the state of the underlying neural activity is unknown since we did not account for the hemodynamic response function (HRF) in our reduced model. Therefore, the input in the model captures the external drivers of regional BOLD and only indirectly, the underlying neural activity. In order to determine the *parameters of the system* (1), i.e., (*A, B*, 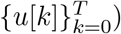), we need to solve an optimization problem that minimizes the distance between the system’s state *x*[*k*] and the estimate of that state given by 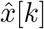 driven by the unknown quantities. Specifically, we have the following optimization problem:

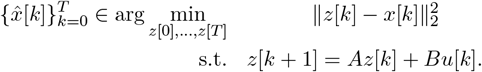

Notice that this problem is more challenging than the usual least squares problem considered when the parameters of the system are known^33^. Thus, similar to the method develop by^26^, we perform the following steps: (i) we assume that the state *z*[0] = *x*[0], and 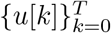 is identically zero, to find an approximation to *A*;(ii) assuming *A* is given by the initial approximation, we provide a sparse low-rank structure to matrix *B* and we find an approximation to both *z*[0] and 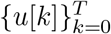, which suffices to obtain *z*[0],…, *z*[*T*] subsequently, 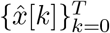; and (iii) assume 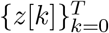 and 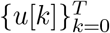 are as approximated in step (ii) and determine an approximation to *B*. The process consists of executing step (ii) and (iii) iteratively. Our experiments reveal that the estimated parameters converge after a few iterations in both synthetic and fMRI time series (Supplementary Fig. 18). Additionally, to force the inputs to be used as little as possible, since otherwise they could contain all the required information to obtain the sequence 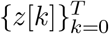 (e.g., consider *A* to be zero and *B* to be the identity matrix), the optimization objective is rather given by 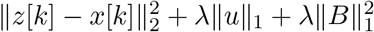, which penalizes the use of the input with a weight *λ* > 0. – See section SI1 for algorithm details.

We will demonstrate in the following results section that unaccounted external inputs result in error in estimation of system matrix *A*. Therefore, in a modified version of this algorithm, in step (i) we estimate *A* from *x′*[*k*] measured during an extended window without external stimulation (e.g., resting-state). Next, we repeat steps (ii) and (iii) iteratively – as detailed above. Since we did not know the true dimensionality of the external inputs, we approximated the dimensions of the input matrix B by performing principal component analysis on the residuals of the models. As seen in Supplementary Fig. 19, principal components 1-25 capture more than 80% of variance in the average residuals and more than average 60% of subject-level residuals’ variance across all tasks. In addition, we compared the goodness-of-fit of the LTI model with and without external inputs using Akaike information criterion (AIC) ^34^. Our results demonstrate that incorporating external inputs does not results in overfitting and improves the model’s fit - an effect most pronounced in higher dimensional input matrices (Supplementary Fig.20). Finally, we demonstrate that we identify the external inputs during the motor task similarly at high-dimensional input matrices (Supplementary Fig. 6), as indicated by the high correlation (>0.8) of inputs estimated using input matrix dimensions higher than 25 (Supplementary Fig. 6I). Therefore, we select *p* = 25 for input matrix B to estimate the inputs from task fMRI time series.

### 2.2 Spectral analysis of an LTI system

Provided an LTI description of the system dynamics (1), the autonomous evolution of the dynamical system can be decomposed in a so-called *eigenmode decomposition*. Briefly, consider the *n* eigenmodes (i.e., eigenvalues and the corresponding eigenvectors) associated with *A*. Each eigenmode corresponds to an eigenvalue-eigenvector pair (*λ_i_, v_i_*) satisfying *Av_i_* = *λ_i_v_i_*, and it describes the oscillatory behavior for a specific direction *v_i_*.

Specifically, for any given eigenvalue *λ_i_* represented in polar coordinates (*θ_i_*, | *λ_i_*|), we have that it captures the *frequency* characterized as

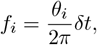

where *δt* corresponds to the sampling frequency, and the *time scale* given by

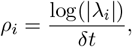

which can be interpreted as the *damping rate*.

In particular, we can re-write *A* = *VλV*^T^, where *V* = [*v*_1_,…, *v_n_*] and *λ* = diag(*λ*_1_,…, *λ_n_*) are the matrices of eigenvectors and eigenvalues. Subsequently, we can apply a change of variable as *z*[*k*] = *V*\**x*[*k*], where *V*\* is the transpose conjugate, which implies that 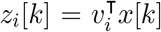 is a weighted combination described by the *i_th_* eigenvector associated with the *i_th_* eigenvalue. Hence, this can be understood as the spatial contributions of the *n* ROIs at a given (spatiotemporal) frequency *f_i_*. Additionally, we can revisit the damping rate of the process in such direction *v_i_* by reasoning as follows: first, we can recursively obtain |*z_i_*[*k*]| = |*λ_i_*|^*t*^|*z_i_*[0]|. Therefore, we have the following three scenarios: (*i*) |*λ_i_*| < 1; (ii) |*λ_i_*| > 1; and (*iii*) |*λ_i_*| = 1. Incase (i) and (ii), we can readily see that |*z_i_*[*k*]| → 0 and |*z_i_*[*k*]| → ∞ as *k* → ∞, respectively. Lastly, in scenario (iii), or practically, when |*λ_i_*| ≈ 1, we have that the process oscillates between *stability* and *instability*, and therefore these dynamics are refer to as *meta-stable*.

In summary, the dynamical process *z*(*k*) describes the spatiotemporal brain BOLD signal evolution. Specifically, the timescales are encoded in the eigenvalues and the spatial contributions of the different ROIs are described by the eigenvectors with a spatiotemporal timescale described by the associated eigenvalues.

### 2.3 Dataset and Preprocessing

We used data from the Human Connectome Project (HCP). As part of the HCP protocol, subjects underwent two separate resting-state scans along with seven task fMRI scans, both of which included two sessions. All data analyzed here came from these scans and was part of the HCP S1200 release. The fMRI protocol (both resting-state and task) includes a multi-band factor of 8, spatial resolution of 2 mm isotropic voxels, and a TR of 0.72 sec (for more details see^35^). Subjects that completed both resting-state scans and all task scans were analyzed. Each of the scanning sessions included both resting-state and task fMRI. First, two 15-minute resting-state scans (eyes open and fixation on a cross-hair) are acquired, for a total of 1 hour of resting-state data over the two-day visit. Second, approximately 30 min of task-fMRI is acquired in each session, including 7 tasks split between the two sessions, for a total of 1 hour of task fMRI (for details see^36^).

The 96 subjects with the lowest mean frame wise displacement were considered in our study, where we utilized a cortical parcellation with *N* = 100 parcels that maximizes the similarity of functional connectivity within each parcel ^37^. We preprocessed resting-state and task data using similar pipelines. For resting-state, the ICA-FIX^38;39^ resting-state data provided by the Human Connectome Project were utilized^40^, which used ICA to remove nuisance and motion signals. For task data, CompCor^41^, with five components from the ventricles and white matter masks, was used to regress out nuisance signals from the time series. In addition, for the task data, the 12 detrended motion estimates provided by the Human Connectome Project were regressed out from the time series. For both task and resting-state, the mean global signal was also removed in an effort to remove the autocorrelated non-physiological noise and reduce the model estimation error^42^.

### 2.4 Statistics

We performed student’s *t*-test and Welch’s *t*-test^43^ to test the statistical significance of the differences between the distributions of interest. Nonparametric Wilcoxon rank-sum test^44^ were utilized for comparisons of distributions with non-normal profiles. We corrected calculated test statistics for multiple comparisons using false discovery rate (FDR) method^45^, as well as the more conservative Bonferroni method^46^. To identify the task-specific fluctuations in the average estimated inputs, for each brain regions we compared task-related inputs to those estimated from resting-state time series (paired *t*–test, *p* < 0.05, FDR). In addition we also generated phase-randomized null time series from each subjects’ BOLD times series for the task time series. Next, for each brain region, we compared the average empirical and null estimated inputs for each time point (paired *t*–test, *p* < 0.05, FDR).

To identify estimated inputs that display changes that correspond to different task conditions in the motor paradigm, we first performed a principal component analysis (PCA) on all estimated inputs (*U*) concatenated over all subjects. Next, we identified a single input with the highest absolute principal component (PC) loading for every component. We then multiplied the selected inputs with negative PC loadings by –1. Next, we separately fitted a multiple linear regression model for each PC’s inputs (*U*) using the known task-regressors. We created task-regressors for different conditions by assigning every sample to baseline (0) or one of six events (i.e., visual cue, left hand, right hand, left foot, and right foot movements) based on their temporal proximity to events’ onsets and offsets. We repeated this analysis by shifting task-regressors by different lags (0–12 TRs) to identify the lag that produces the best fit (i.e., highest *R*^2^ values) for each region. Finally, we performed *t*–tests on estimated coefficients at the group-level to identify task conditions similarly echoed in estimated inputs associated with each PC across participants. We also identified brain regions that correspond to the identified inputs by performing group-level region-wise *t*–tests on input matrix B elements that correspond to inputs *U* identified by PCs.

We examined the estimated inputs’ non-stationarity using two methods. First, we used a sliding window approach to examine temporal fluctuations of estimated inputs’ means over resting-state scans for all brain regions, measured from the windowed-means’ standard deviation. Second, we used the nonlinear non-stationarity index introduced by^47^, with *α* = 0.9 and *β* = 1 exponent parameters following their study, where *α* and *β* parameters control the relative weighting between the importance of long versus large excursions in time series. Therefore, non-stationarity indexes with our selected parameters give marginally greater weighting to excursions’ height. Finally, to test the group-level significance of both non-stationarity metrics, we first normalized the values across all brain regions. Next, we used the *t*–test (FDR corrected for multiple comparisons across all brain regions) to establish the statistical significance of the measured non-stationarities across patients. Traditionally, researchers have commonly used the 0.05 as the statistical significance level, though the choice is largely subjective. Therefore to convey the probabilistic nature of the statistical analysis and the proper interpretation of statistical test results, in the manuscript, we refer to results of the commonly accepted statistical threshold of 0.05 as “significant” and the more conservative thresholds of 0.0005 or lower as “highly significant”.

## 3 Results

### 3.1 Retrieving the external inputs to a synthetic LTI system

We use the proposed method to explicitly model the contributions of internal system dynamics and external inputs on the BOLD signal during rest and task. To build intuition, we begin by estimating the internal system parameters and unknown inputs using data simulated from a synthetic LTI model (Eq. 1) with four states representing four brain regions. We first simulate the dynamics of our model (Fig. 1A), where each region is driven by random internal noise, and only one region is driven by an additional square pulse train (Fig. 1B). For details regarding the simulation see the Supplementary Information (SI) section SI2. Next, we estimate internal system parameters (4 × 4 matrix of interactions) and unknown inputs from the simulated time series, and to recover the spatial and temporal profiles of the pulse train input (Fig. 1C). Although the estimated inputs (green line) fluctuate time-locked to the ground-truth input, their temporal profiles notably differ. We hypothesize that this divergence arises from the error in system matrices estimated during periods with external stimulations. In Fig. 1D, we show that the LTI system parameters receiving time-varying external inputs can falsely appear to change and diverge farther from the ground-truth when examined over periods with external stimulation.

**Figure 1:**
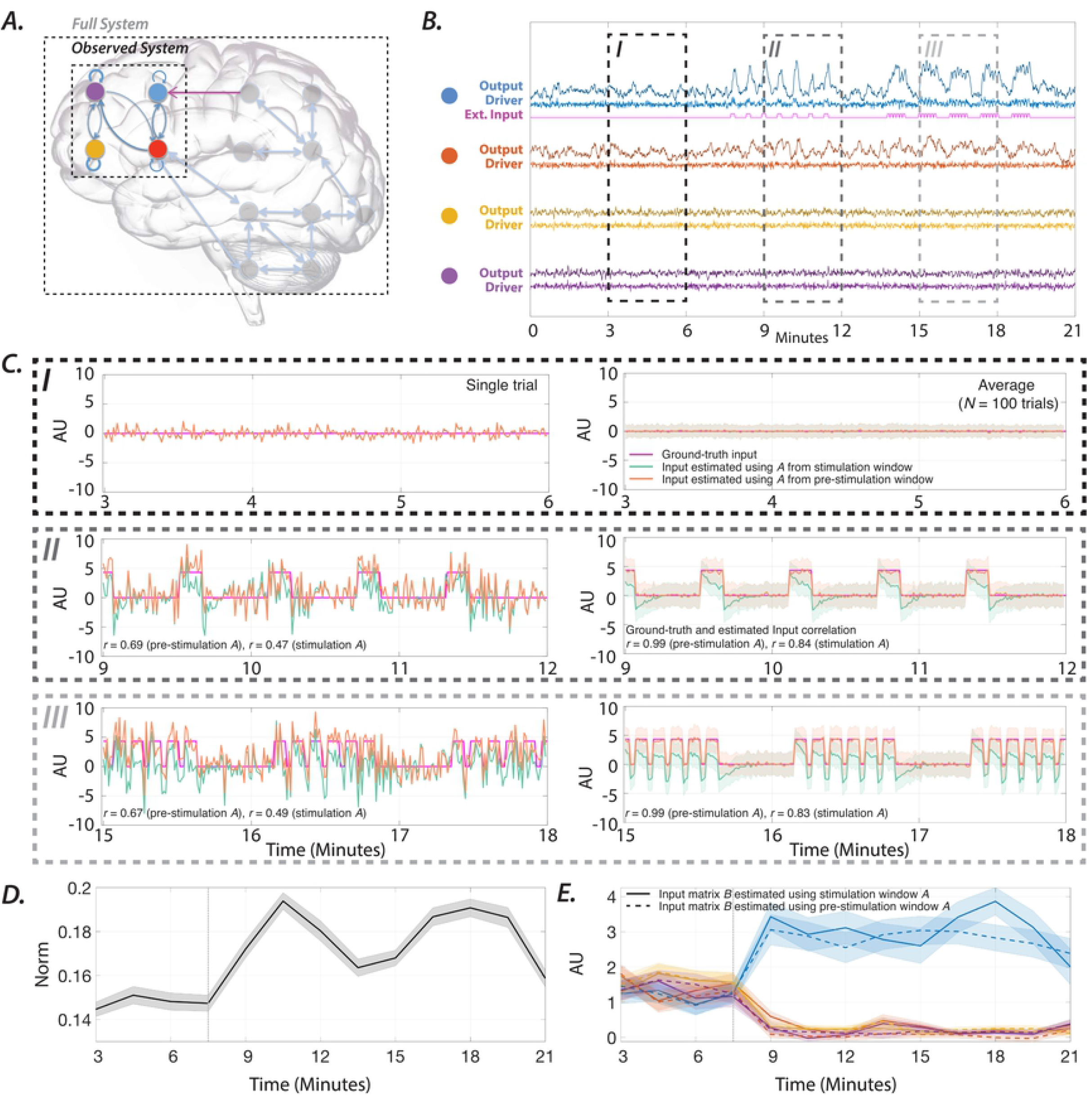
Synthetic LTI system with unknown inputs. ***(A) A*** schematic of the brain as a network, where the nodes represent brain regions, and the edges represent connections between regions. The activity of four *observed* regions is modeled as a four-dimensional LTI system, and the influence of the unobserved regions and external stimuli into each node as an unknown driver. The synthetic system matrix is designed with eigenmodes oscillating at 0.01 and 0.06 Hz to mimic the frequencies of BOLD signal’s neurophysiological component. ***(B)*** Simulated time-evolution of each node’s activity (sampling rate = 1.4 Hz) is color-coded and shown in the presence of drivers, namely the internal noise and the external input (brighter colors). Only the blue node receives external input indicated by the magenta line. Three periods (I–III) are highlighted dashed lines. At period I (3–6 min), there is no external stimulation. At period II (9–12 min), the blue node is stimulated in 25 samples = 18 seconds blocks, interleaved with similarly sized rest periods. At period III (15–18 min), the blue node is stimulated for 7 samples = 5.04 seconds, with inter-stimulus intervals of 3 samples = 2.16 seconds. ***(C)*** Left panels show the estimated inputs to the blue node (green line, arbitrary units AU) estimated from a single simulation. The panels on the right show the average input and its standard error over 100 simulations. ***(D)*** The average 2-norm and standard error of the difference between the system’s true and estimated matrices of a 3-minute sliding window. ***(E)*** The color-coded lines show the average (and standard error) loading of each node on input matrix *B*.

Consequently, we hypothesize that system matrices estimated from periods without external inputs would improve our ability to capture the unknown inputs’ profile accurately. Fig. 1C shows that using a fixed system matrix estimated from periods without external inputs significantly increases the similarity (correlation) to the ground-truth inputs. We also demonstrate that although estimated inputs contain noise, averaging inputs estimated over 100 simulations results in highly accurate estimations (correlation = 0.99). The significant (Wilcoxon rank-sum test, Bonferroni *p* < 0.0001) changes in the input matrix *B*’s loading for estimation windows overlapping the external stimulation periods, reveals the unknown external inputs’ spatial profile (i.e., the blue input node) (Fig. 1E). Together, these results demonstrate that external inputs can increase estimation error in system matrices, and consequently, input parameters. More importantly, these results also show that identifying system matrices from periods without external stimulation allow an accurate estimation of unknown external inputs’ spatiotemporal profiles.

Next, we generate synthetic time series by stimulating LTI systems, parameters of which were estimated from subjects’ resting-state BOLD time series. We set external inputs’ magnitude such that the global average stimulus-induced changes in normalized simulated outputs match the largest average task-related changes in a sample (social) task. We confirm that similar to the low-dimensional example in Fig. 1, our approach is able to extract synthetic external inputs to high-dimensional LTI models of BOLD signal dynamics (Fig. 2). Likewise, employing system parameters estimated from periods without external stimulation results in a significant (*t*–test, *p* < 0.05, *p* = 6.6 × 10^-65^ and *p* = 2.9 × 10^-66^ for 1000 TR- and 250 TR-long estimation windows, respectively) increase in the similarity between the ground-truth and estimated inputs (Fig. 2F). The notably higher similarity between the average estimated to ground-truth inputs than that of subject-level estimated inputs suggests that profiles of external inputs are correctly approximated although with noise. Together these results demonstrate the utility of our framework in identifying external inputs to LTI systems, and highlight the importance of accurate estimation of model parameters.

**Figure 2:**
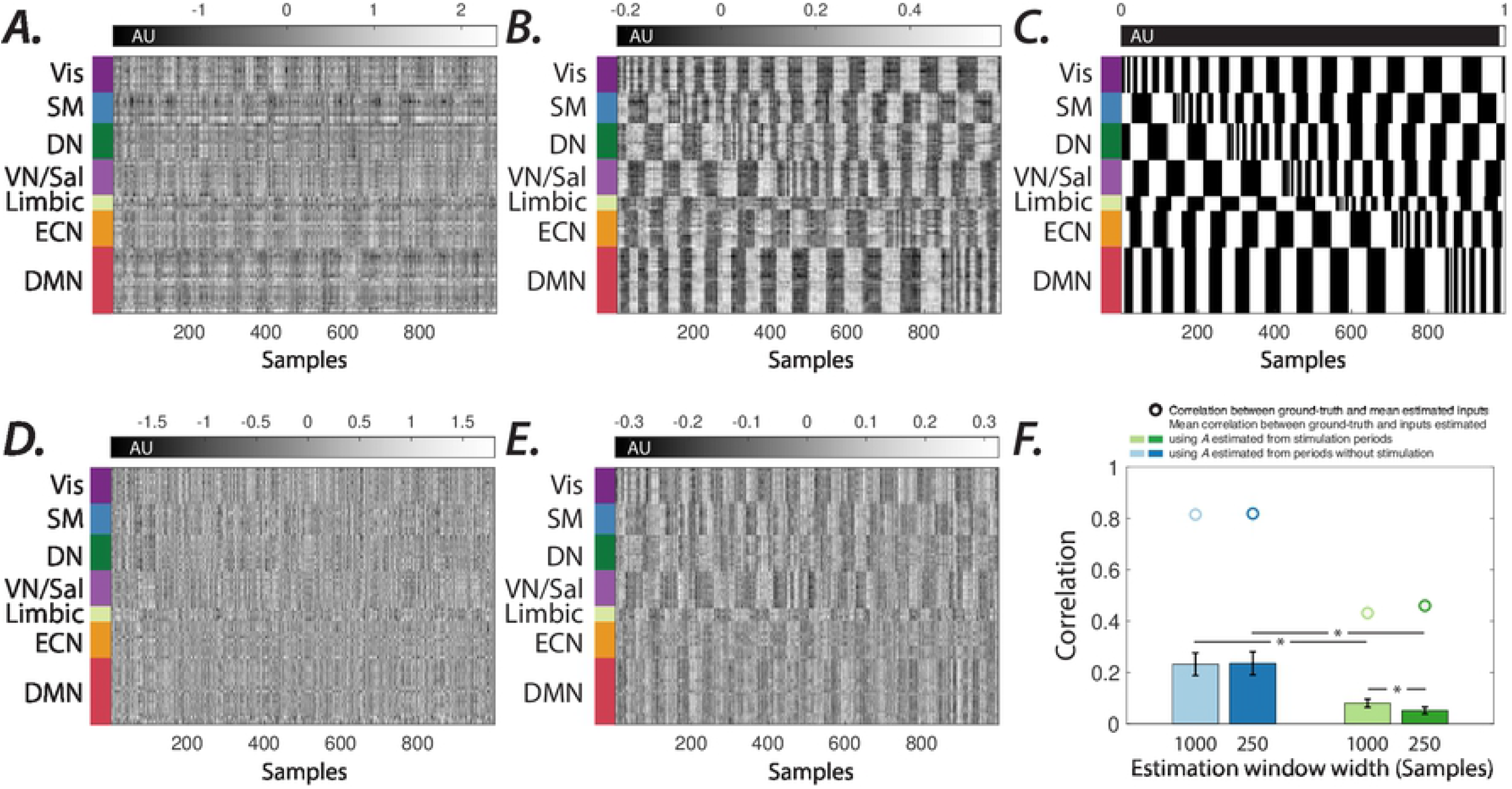
Extracting spatiotemporal profiles of unknown external drivers in simulated brain dynamics. ***(A&D)*** Estimated external inputs (i.e., *B* × *U*) to all brain regions from synthetic time series generated from a sample subject’s internal system parameters and (***B&E***) the average estimated external inputs across all subjects (input matrix *B* dimension = 7, regularization factor = 0.5). Brain regions (y-axis) are sorted based on resting-state networks identified by^48^, namely the visual (Vis), sensory/motor (SM), dorsal attention (DN), ventral attention/salience (VN/Sal), limbic, executive control (ECN), and default mode network (DMN). System parameters in panels ***D&E*** are estimated from the stimulation window, however system parameters in panels ***A&B*** are estimated from same-length windows without external inputs. ***(C)*** Ground-truth synthetic inputs over 1000 samples (TR = 0.72 sec). ***(F)*** The similarity between ground-truth and estimated inputs. The system matrix *A* estimated from windows without external stimulation results in a significantly higher correlation between the vectorized estimated external and ground-truth input matrices (*t*-test, *p* < 0.05, *p* = 6.6 × 10^-65^ and *p* = 2.9 × 10^-66^ for estimation windows with 1000 and 250 samples, respectively), compared to system matrix *A* estimated from the stimulation windows (indicated by ‘*’ markers). The smaller estimation windows significantly (*t*-test, *p* < 0.05, p = 1.15 × 10^-45^) reduce the estimated and ground-truth inputs’ similarity, only for the system matrix *A* estimated over stimulation windows (indicated by ‘*’ markers). The correlation values between the ground-truth and group average estimated inputs are indicated by ‘o’ markers.

So far, we have examined the LTI system’s response in a low recording noise level (signal-to-recording noise =1000). Next, we examine the accuracy of the retrieved model and input parameters at different recording and internal noise levels. The contributions of the recording and internal noise to the BOLD signal, for the most part, are unknown quantities. However, they play an essential role in our ability to capture external inputs accurately. Simulating the system’s response magnitude and variance (i.e., *t*-values) at various recording and internal noise levels show how different noise levels can lead to seemingly similar outputs.

Moreover, at high noise levels, the error increases notably in the system parameters estimated from periods without external inputs, and consequently, in the estimated input parameters during stimulation periods. Interestingly, at such high noise levels, the system matrices estimated during stimulation periods more accurately recover external inputs than those estimated during periods without stimulation (Supplementary Fig. 1). These observations suggest that the choice of system matrices and the goodness-of-fit of the estimated inputs can further provide insight into the empirical noise levels. In the following, we consider the proposed methodology in the context of quantifying important spatial and temporal features of the internal system dynamics and external inputs estimated from the HCP resting-state and task fMRI scans.

### 3.2 Capturing external drivers of BOLD signal

#### 3.2.1 Brain’s large-scale oscillatory modes display heterogeneous spatiotemporal profiles

We begin by showing that the estimated system parameters during restingstate reliably capture and reproduce known brain functional organization. Further, because these parameters reside within a quantitative dynamical model, we simultaneously capture both spatial (regions that are co-active) and temporal (oscillation frequency) information through the *eigenmodes* of our estimated system. Specifically, each eigenvector indicates an independent pattern of co-active regions, and its corresponding eigenvalue determines both the oscillation frequency and the change in amplitude of the activation patterns. Intuitively, if we initialize our estimated system state to a pattern of activity corresponding to an eigenvector, then the system states would oscillate and dampen according to the associated eigenvalue’s characteristics (see more details in Materials and Methods section).

To capture the spatial and temporal patterns of activity, we use our method to estimate the internal system parameters from the resting-state time series (1200 TR ≈ 14.5 min). The high stability of the (i.e., slow damping rate) low-frequency eigenvalues as seen in Fig. 4A indicates that the system’s outputs are dominated by lower frequency oscillations. To identify the eigenmodes with similar spatial patterns across subjects, we aggregate all subjects’ eigenvectors and perform *k*-means clustering analysis. We provide the course (*k* = 2) and finer scale (*k* = 4) clusters in Supplementary Fig. 2.A and Fig. 3, respectively (for details regarding clustering method see SI5 and Supplementary Fig. 3). To test the spatial inhomogeneity in the frequency and damping of these clustered eigenvectors, we performed a pairwise comparison between the distribution of eigenvalues corresponding to the eigenvectors in each of the clusters (bootstrap *n* = 50,000, Bonferroni corrected *p* < 0.05). We found significant differences in the frequencies and damping rates between all cluster pairs, except for the comparison between the frequencies in clusters 3 and 4).

**Figure 3:**
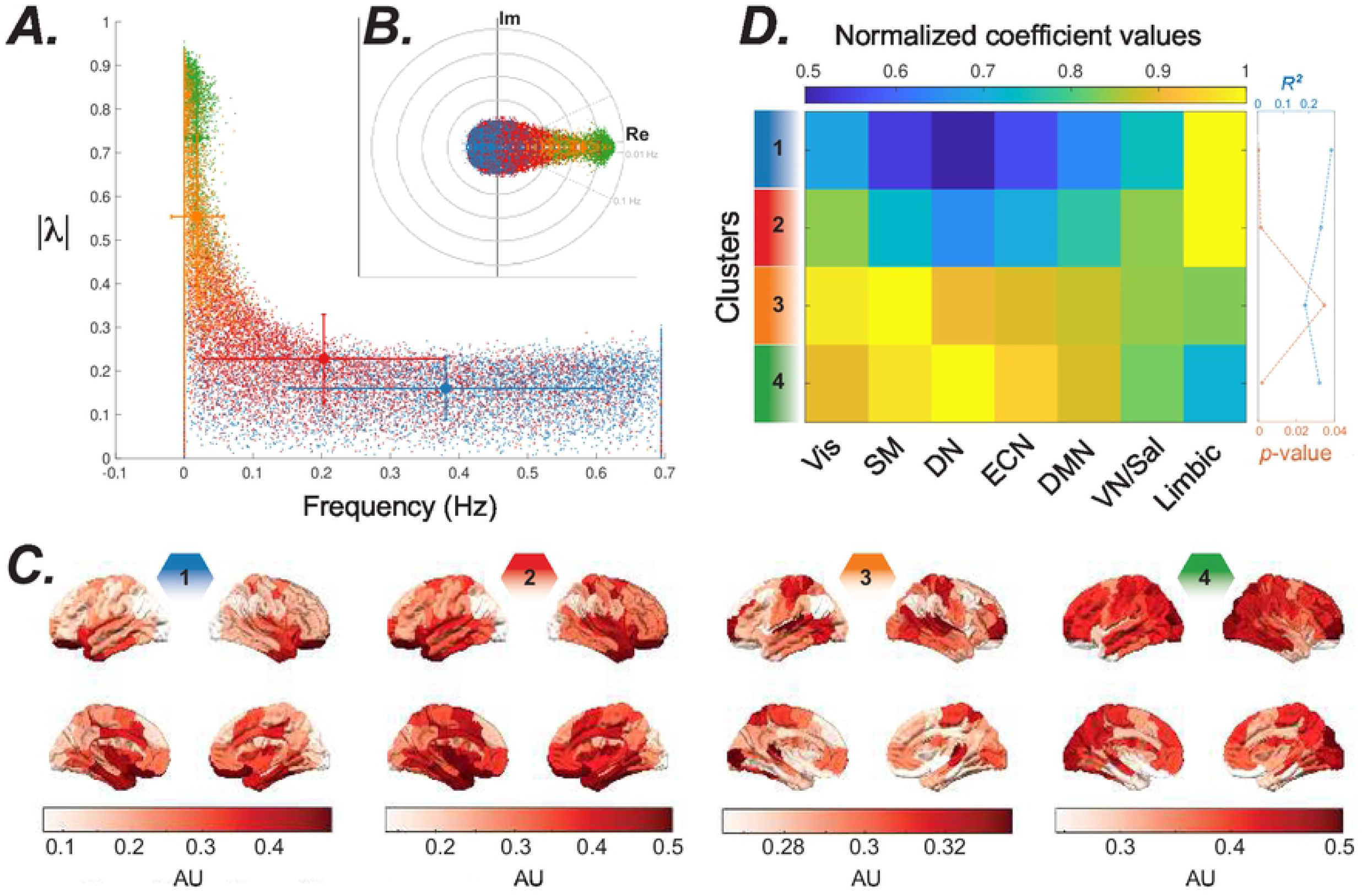
Eigenmodes estimated from the full (1200 TR ≈ 14.5 min) restingstate time series. ***(A)*** Distribution of frequency versus stability of eigenvalues during resting-state. Clustering the eigenvalues based on their eigenvector’s similarity highlights the spectral profile of different systems. All eigenvectors from all subjects were normalized and grouped into 4 clusters using the fc-means clustering algorithm. We color-coded the clusters identified across subjects and all resting-state sessions (*n* = 4). ***(B)*** the inset plot shows the eigenvalues’ distribution. ***(C)*** The brain overlays represent the spatial distribution of the eigenvector associated with an eigenvalue (displayed with the same color code) that is at the centroid of each cluster. ***(D)*** The similarity between eigenvector clusters’ centroids and the resting-state networks. We performed spatial multiple linear regression analyses using all resting-state networks identified by^48^, namely the visual (Vis), sensory/motor (SM), dorsal attention (DN), ventral attention/salience (VN/Sal), limbic, executive control (ECN), and default mode network (DMN) as the explanatory variables, to show which resting-state networks overlap with the eigenvector clusters’ centroids shown in the panel. The color-coded matrix shows the estimated normalized (divided by maximum value at each row) coefficients of the regression, calculated separately for every eigenvectors’ cluster’s centroid. The plot on the right shows the *p*-value and *R*^2^ calculated for each cluster centroid.

**Figure 4:**
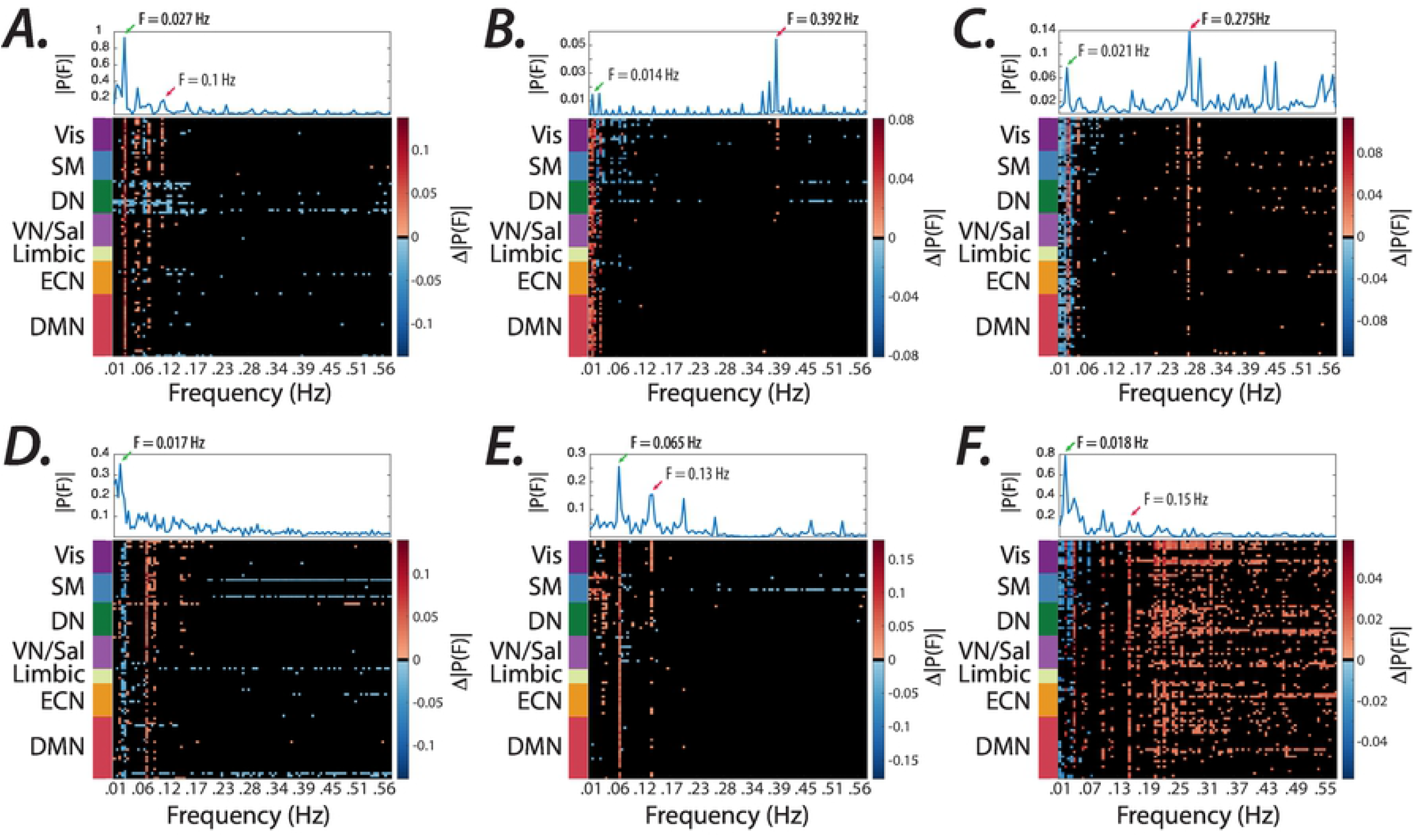
Matching the spectral profile of the known and estimated external inputs. ***(A-F)*** The difference between the average Fourier transform of the estimated inputs to all brain regions during tasks compared to that of other task conditions (see Materials and Methods for details). Top panels display the average (two sessions) spectral profile of the known boxcar regressors for each task (see Supplementary Fig. 4). Note the significant changes in the spectrum at expected task-specific frequency peaks across several brain regions, at low (< 0.1 Hz) and high (> 0.1 Hz) frequencies represented with red and green arrows, respectively. Frequencies for which brain regions did not pass the significance level (Wilcoxon rank-sum test, FDR p < 0.0005) are represented in black.

#### 3.2.2 Task-specific increases in the extra-cortical input’s power

Up to now, we provided evidence that the system dynamics can capture the spatial and temporal behavior of resting-state brain networks. Next, we try to assess if the task-induced dynamics are driven by the external inputs, retrieved by the proposed method. The sensory inputs to the brain are some of the major drivers of cortical dynamics. Therefore, we hypothesize that the external inputs to the subjects’ brains, as estimated by the proposed method, will mirror real-time changes present in these task regressors (see Supplementary Fig. 4 for details regarding the task regressors).

To test this hypothesis, we apply our method to the fMRI activity to estimate the internal system parameters and external inputs for each subject during task performance (i.e., social, gambling, motor, working memory, language, and relational). Then, we compare the average estimated inputs’ frequency spectrum for each task. Statistical tests (Wilcoxon rank-sum test, FDR corrected, *p* < 0.0005) reveal highly significant unique peaks, matching the expected external task-specific frequencies (Fig 4). Note that the distinct task-induced peaks are identified at low (< 0.1 Hz) and high (> 0.1 Hz) frequencies, even as high as 0.3–0.4 Hz (Fig 4B-C.)

#### 3.2.3 Task-specific profiles of extra-cortical inputs

Next, we consider an LTI framework to quantify spatial and temporal features of external inputs to the brain using HCP’s motor task dataset. The motor task comprises 3-second long visual cues, where participants are asked to either tap left or right fingers, squeeze left or right toes, or move their tongue over 12-second long periods following the visual cue’s offset. We select the motor task since the high dimensionality of input and various task conditions in this paradigm allows us to evaluate our framework’s ability to estimate external inputs’ complex spatiotemporal structure. We aim to assess if we can retrieve the external inputs that drive task-induced dynamics. We hypothesize that subjects’ estimated external inputs will mirror realtime changes present in known task regressors. Moreover, due to relatively lower levels of structured external stimulations during resting-state scans, we hypothesize that the system parameters estimated from subjects’ fulllength resting-state time series will increase the accuracy of external inputs estimated from motor task datasets.

Fig. 5 demonstrates estimated inputs (input matrix *B* dimensions = 25, regularization factor = 0.5) to all brain regions (i.e., *B* × *U*) averaged across all subjects during the motor task. These results highlight the brain-wide significant task-specific changes in the estimated inputs when system parameters are estimated from the resting-state time series Fig. 5A-B. We provide evidence of the robustness of these results to changes in the regularization parameter (Supplementary Fig. 5), and to increase the input matrix *B*’s dimension (Supplementary Fig. 6). Conversely, the identified inputs using the system parameters estimated from the subjects’ motor task time series notably reduces our ability to capture the task-related changes (Fig. 5C-D).

**Figure 5:**
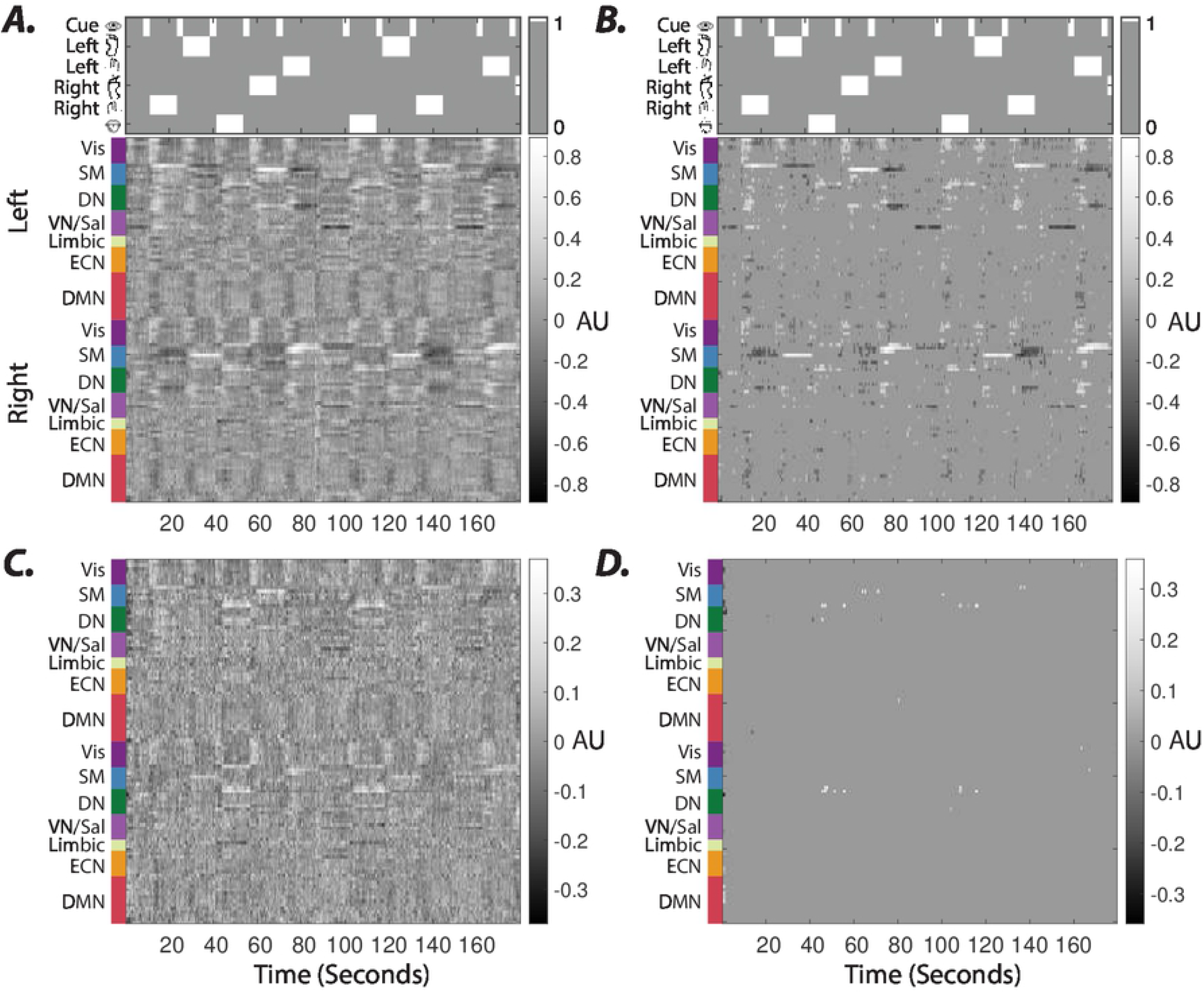
Average estimated external inputs in the motor task. Internal system parameters (i.e., *A* matrices) during full-length resting-state and motor scans were used to estimate the external inputs in panels ***A*** and ***C***, respectively. Panels ***B*** and ***D*** show time points form panels ***A*** and ***C*** with significantly higher or lower average inputs estimated during motor task than resting-state scans (paired *t*–test, *p* < 0.05, false discovery rate (FDR) corrected for multiple comparisons). Top plots in panels ***A&B*** show onsets and durations of visual cues and motor task conditions – left foot, left hand, right foot, right hand, and tongue movement blocks.

We establish these observations’ statistical significance by comparing the external inputs estimated from task datasets against those from subjects’ resting-state scans (paired *t*–test, *p* < 0.05, FDR corrected for multiple comparisons). Comparisons against the phase-randomized null time series also provide converging observations (Supplementary Fig. 7). We also use multiple linear regression analyses to assess the estimated inputs’ similarity to the known temporal profile of the task regressors. Our results demonstrate that external inputs estimated using the full-length resting-state system parameters result in significantly (paired *t*–test, *p* < 0.05, Bonferroni corrected for multiple comparisons) improved fit (measured by *R*^2^ values), compared to system parameters estimated from the motor task (Supplementary Fig. 8). We also find similar results when resting-state system parameters were estimated from a short (250 sample) window that match task scans’ length (Supplementary Fig. 8B). Together these results highlight the importance of the modeled system’s accuracy in capturing a reliable picture of the brain’s external inputs.

Next, we examine the temporal (i.e., *U* matrix) and the spatial (i.e., input matrix *B*) profiles of the external inputs (estimated using resting-state system parameters), to demonstrate how the estimated inputs reveal the dimensionality and the spatiotemporal dependencies of the task-related inputs. Prior works using univariate and multivariate analyses of HCP task datasets have demonstrated that activation induced by the hand, foot, and tongue movements can be localized over the somatomotor network. Therefore, we expect the dimensionality of the external inputs to roughly match or exceed those of task conditions (i.e., six dimensions). As mentioned in the Materials and Methods section, the principal component analysis reveals that in all HCP task conditions, principal components (PCs) 1–25 explain more than 80 % of the variance in the model’s average residuals. Therefore, we choose *p* = 25 as the input matrix *B* dimension in Fig. 6.

**Figure 6:**
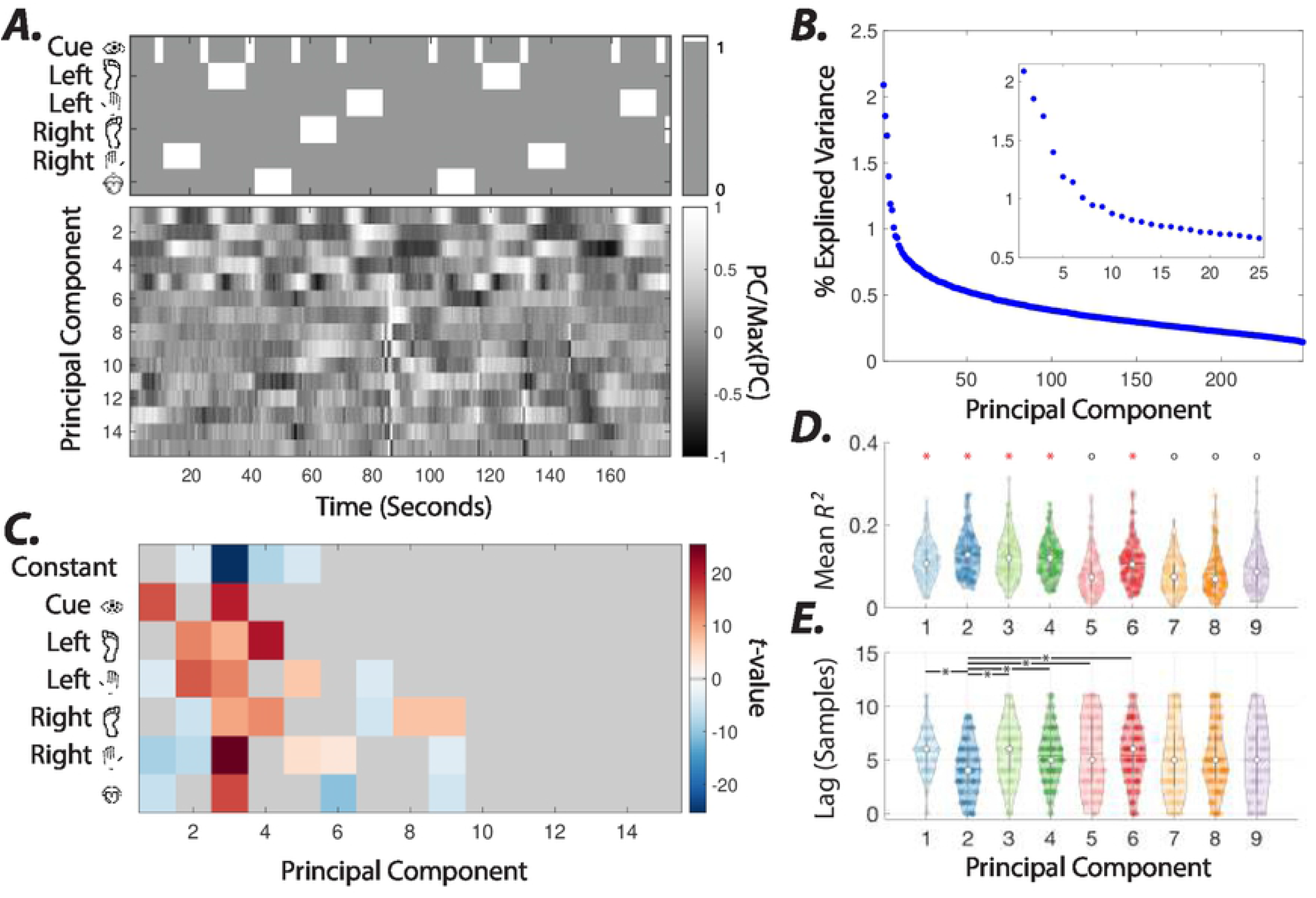
Principal component analysis of estimated external inputs. ***(A)*** Group-level principal components (PCs) 1–15 calculated from concatenated estimated inputs (input matrix *B* dimension = 25) across all subjects. Top plots show onsets and durations of visual cues and motor task conditions. ***(B)*** Percent variance explained by PCs. Insets depict the percent variance explained by PCs 1–25. ***(C)*** t-values calculated from coefficients of multiple linear regression models of estimated external inputs associated with each PC (see methods for details). The average coefficients that fail to pass the significance-level across subjects (*t*-test, *p* < 0.05, Bonferroni corrected for multiple comparisons) are depicted in gray. ***(D)*** Distributions of *R*^2^ values of multiple linear regression models in panel ***B*** for components with significant coefficients. White circles and color-coded horizontal bars indicate the medians and means of distributions, respectively. Pairwise comparison (Wilcoxon rank-sum test, *p* < 0.05, FDR corrected for multiple comparisons) between distributions reveal that *R*^2^ values for principal components marked by red ‘*’ are significantly higher than those marked by black ‘o’ (except for the non-significant difference between PC 1 and PC 9). ***(E)*** Distributions of the number of lags (samples) that results in best fit (i.e., maximum *R*^2^) for PCs 1-9. We used the mean (round to nearest integer) of optimal subject-level lags for analysis in panels ***C*** and ***D***.

We performed principal component analysis on external inputs estimated temporal profiles (i.e., *U*) concatenated across all subjects to identify the input patterns similarly identified over the group. Fig. 6A shows the temporal profile of the concatenated inputs’ PCs 1–15. As seen in Fig. 6B, the first few PCs (≈ 9) explain a relatively larger portion of the variance. Fig 6A shows the high similarity between known task regressors and PCs’ temporal profiles. We quantify this similarity using subject-level multiple linear regression analysis of the estimated inputs using the known task (motor) regressors. We note apparent time lags between the known and estimated inputs. Therefore, we perform the multiple linear regression analysis using various lags. Fig 6D shows distributions of lags (samples) that yield the highest *R*^2^ values for PCs 1–9. Fig. 6B shows the group average coefficients estimated from external inputs associated with each PC (i.e., external inputs with highest PC weights). We used the group average optimal lag (based on *R*^2^ values) identified in Fig. 6E in Fig. 6B. Estimated coefficients have significant values, only in PCs 1—9. These results demonstrate that the estimated inputs provide insight into the extra-cortical drivers’ dimensionality.

Next, we examine spatiotemporal profiles of subject-level estimated inputs associated with these components to understand their relationship to the external stimuli. Fig. 6D demonstrate that compared to other PCs, the inputs associated with PCs 1-4 and 6 fit task regressors relatively better, indicated by significantly (Wilcoxon rank-sum test, *p* < 0.05, FDR corrected for multiple comparisons) higher *R*^2^ values. Fig. 6C reveals that PCs 1-4 and 6 are associated with the visual cue, hand and feet movements (maximum coefficient in left hand), all movements (maximum coefficient in right hand), feet movements (maximum coefficient in left foot), and tongue movements, respectively. Supplementary Fig. 9 shows that the brain regions with the highest average absolute input matrix B values corresponding to PCs 2, 4, and 6 reveal the same regions identified in the somatomotor cortices using general linear model analysis of BOLD time series for hand, foot, and tongue movements.

The input matrix *B* also captures the spatiotemporal relationship between the inputs across different conditions. For instance, Supplementary Fig. 9A shows that hand or feet movements are associated with simultaneous positive and negative (e.g., inhibition or deactivation) inputs to the contra- and ipsilateral somatomotor cortices, respectively. Fig. 7 also shows that PCs 5, 1, and 3 reveal the temporal order of inputs to visual, dorsal attention, and finally, somatomotor cortices following the onset of visual cue. Note that the spatial and temporal profile of PC 5 demonstrates the inverse relationship between inputs to visual and somatomotor cortices. This unexpected temporal profile contributes to the low similarity of PC 5 to task regressors in Fig. 6D. We show that changing the delay between estimated inputs and task regressors changes the coefficient patterns with significant loading (Supplementary Fig. 10). These results demonstrate an early positive relationship of PC 5 input with visual cue blocks, followed by a later positive (negative) relationship with left-hand movements (visual) blocks.

**Figure 7:**
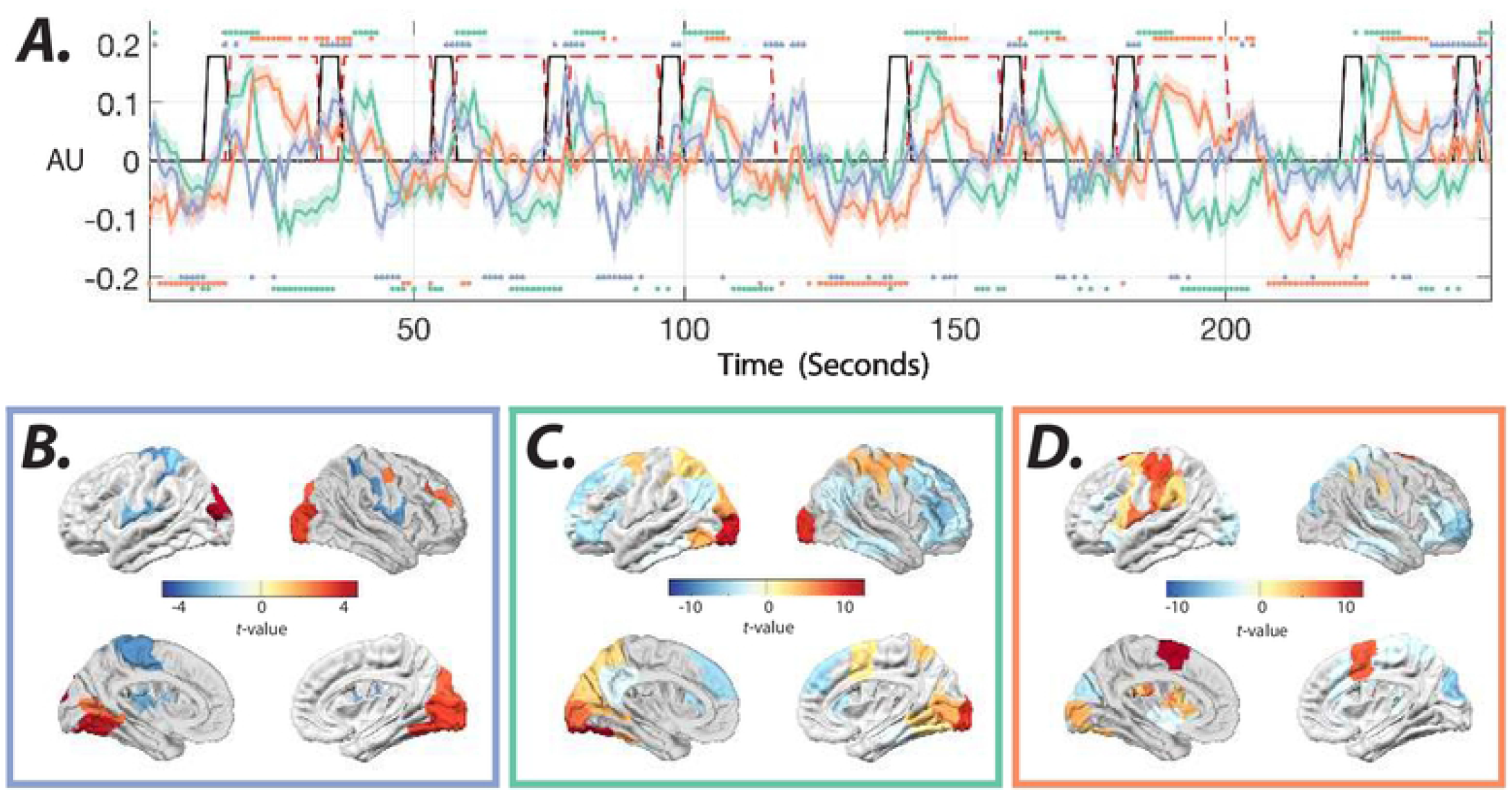
Temporal and spatial profiles of estimated external inputs associated with visual cues. ***(A)*** Color-coded lines show the mean and standard error (shaded area) of estimated inputs with the highest subject-level loadings for PCs 1 (green), 3 (orange), and 5 (blue). Time points with significant (*t*–test, *p* < 0.05, FDR corrected for multiple comparisons across time points) divergence from zero are marked with color-coded dots. The black and dashed red lines show the visual cue and motor task blocks, respectively. Color-coded panels ***(B-D)*** show the *t*–test values of brain regions with significant (*t*–test, *p* < 0.05, FDR corrected for multiple comparisons across ROIs) loadings on input matrix B rows corresponding to the aforementioned PCs.

Finally, in Fig. 6C we demonstrate that PCs 7, 8, and 9 are primarily associated with the right foot movement blocks. However, the significantly smaller *R*^2^ values of these PCs than other PCs in Fig. 6C indicates the lower similarity of corresponding estimated inputs’ temporal profiles to those of task regressors. Closer examination of these inputs’ spatiotemporal profiles reveals that in addition to changes related to left-hand movements, these PCs capture the rapid sequence of inputs to frontal and somatomotor cortices following the motor task block’s offset and the baseline (i.e., no task) onset (Supplementary Fig. 11). Together, these results suggest that an LTI model of cortical dynamics can reveal the unknown spatiotemporal profiles of the BOLD signal’s external task-related drivers.

We provide additional analysis and discussion on model parameters and their effect on the reported results in the Supplementary Information (SI) document. We explored sparsity constraints on the system and input parameters in SI5. Supplementary Fig. 12 demonstrates that increasing the system matrices’ sparsity reduces the model’s goodness-of-fit (measured using the AIC criterion). In the same vein, the increased spatiotemporal sparsity of the inputs overall reduces the accuracy (measured using the *R*^2^ value of the linear regression) of the estimated inputs (Supplementary Fig. 13). Nevertheless, estimated inputs’ group-level PCA reveals that the higher sparsity constraints can improve the accuracy of specific empirically identified input patterns (Supplementary Fig. 14). In addition, we examined the effect of the estimation window’s size on the input’s accuracy in SI6. These results show that a smaller estimation window (3 min) provide comparable results to the full-length window, however overall it increases the accuracy of mean inputs to many brain regions (Supplementary Fig. 8) and several main input patterns (Supplementary Fig. 15). Finally, we explored the sensitivity of the identified input patterns to the factorizations method in the SI7. These results demonstrate that PCA decomposition of the model’s residuals reveals the analogous primary input patterns (Supplementary Fig. 16) uncovered by our spatiotemporal regularization scheme.

#### 3.2.4 Non-stationarity of inputs to resting-state networks

So far, we showed that adopting a time-invariant model of the intrinsic relationship between large-scale brain regions allows us to extract the unknown external drivers of cortical dynamics. Our results demonstrate that the resting-state paradigm serves as a viable option for a more accurate estimation of internal system parameters. However, sensory and other extra-cortical inputs are still present during resting-state scans, resulting in system parameters and input estimation errors. Despite the estimation error in the external inputs’ profile, we hypothesize that quantifying the non-stationarity of the estimated resting-state inputs provides information on the external factors that contribute to resting-state BOLD signal non-stationarities.

We quantify estimated inputs’ non-stationarity for every brain region (i.e., *B* × *U*) from the temporal fluctuations (i.e., standard deviation) of external inputs’ means, measured using a sliding window. Fig. 8 shows brain regions that exhibit significantly high input means’ fluctuation across different sliding window sizes (see methods for details). We demonstrate the results for sliding windows of 6, 24, and 50 samples (TR = 0.72 sec) lengths and half window-length shifts. We also measure the non-stationary of external inputs during resting-state scans using the nonlinear measure developed by^47^ and find converging results (Fig. 8B). We find several brain regions within DMN consistently display high non-stationarity values. Statistical comparisons between the quantified non-stationarity of estimated inputs to identified brain regions in Fig. 8 reveal the significantly (Welch’s *t*-test, *p* < 0.05, Bonferroni corrected for multiple comparisons) higher non-stationarity of external inputs to identified DMN regions relative to several other resting-state networks (Supplementary Fig. 17). Together, these results reveal that time-varying external inputs may partly contribute to the previously reported resting-state BOLD signal’s non-stationary, and the LTI model offers an avenue to determine the spatiotemporal profiles of these unknown external sources.

**Figure 8:**
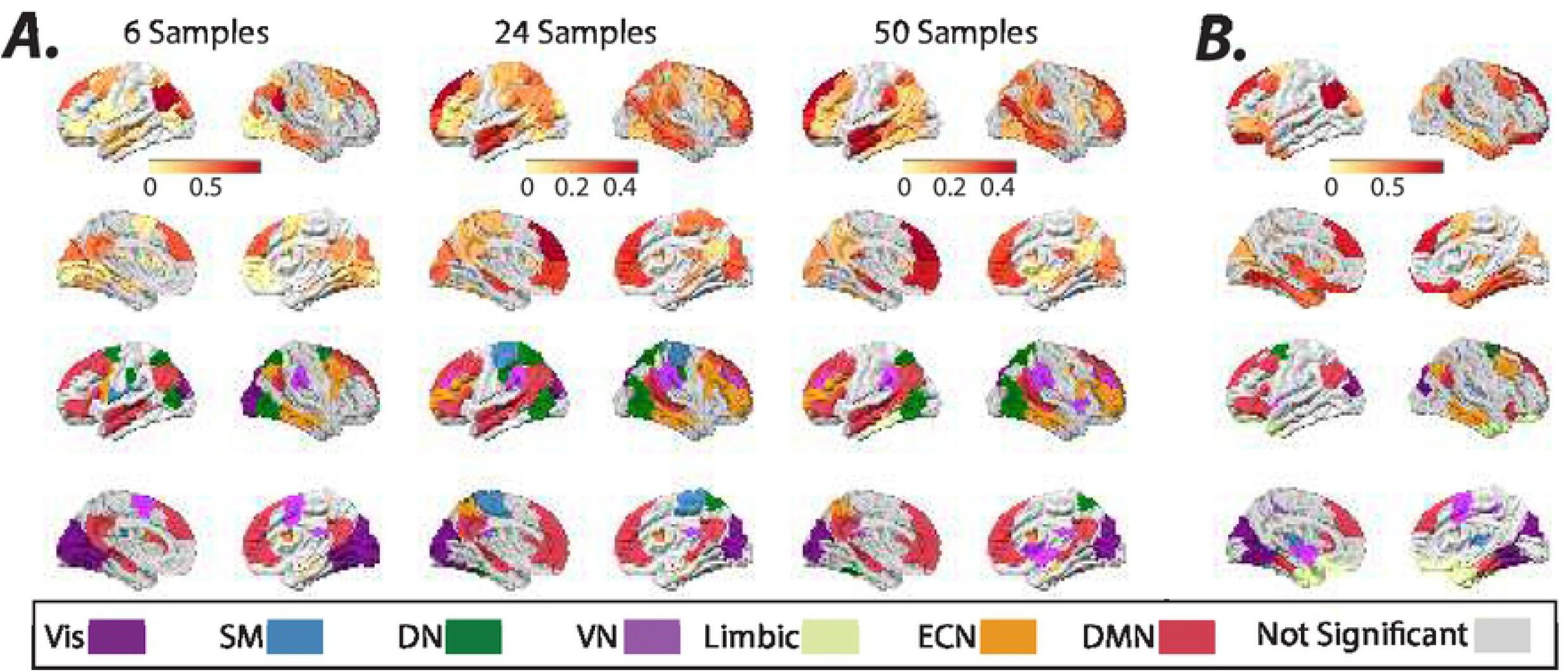
Non-stationarity of estimated external inputs over resting-state scans. ***(A)*** Brain overlays on top panels highlight regions with significantly (*t*–test, *p* < 0.05, FDR corrected for multiple comparisons) high normalized (z-scored over all brain regions) fluctuations (i.e., standard deviation) in the normalized (z-score) estimated inputs’ means, measured using sliding windows (6, 24, and 50 samples window length, TR = 0.72 sec). ***(B)*** Brain overlays on top panels highlight regions with significantly (*t*–test, *p* < 0.05, FDR corrected for multiple comparisons) high normalized (z-scored over all brain regions) nonlinear non-stationarity index developed by^47^, calculated from the normalized (z-score) estimated inputs. The color-coded regions in the bottom plots in panels ***A*** and ***B*** highlight the allegiance of brain regions in top panels to the seven resting-state networks identified by^48^.

## 4 Discussion

Based on the theory of embodied cognition, the evolution and emergent function of the brain can be best understood in the context of the body and its interactions with the environment^49;50;51;52^. In this view, the information does not exist in an abstract form outside the agent, instead, it is actively created through the agent’s physical interaction with the environment^52^. Therefore, understanding the native structure of the external inputs to the brain, as well as the interaction between the brain and its exogenous drivers, is germane to understanding the functional dynamics of the embodied brain^53^.

What are the external drivers of BOLD signal? Current theories suggest that cortical outputs reflect changes in the balance between the strong recurrent local excitation and inhibition connectivity, rather than a feedforward integration of weak subcortical inputs^54^. Changes in this balance heavily affects the local metabolic energy demands and consequently the regulation of cerebral blood flow and the BOLD signal, despite the net excitatory or inhibitory output of the circuits^55^. Inhibition in principle can lead to both increases^55^ and decreases^56;57;58^ in metabolic demands^59^. Moreover, cortical afferents and microcircuits can function as *drivers* by transmitting information about the stimuli, or alternatively as *modulators* by modulating the sensitivity and context-specificity of the response^60;61;62^. Excitatory sensory information, transmitted mostly via glutamatergic or aspartergic drivers, combined with the strong evoked recurrent GABAergic interneurons are a major part of neurotransmission dynamics, which in turn affect the local cerebral blood flow (CBF) ^55^. Likewise, regulation of cortical excitability mediated by neuromodulatory neurotransmitters including acetylcholine^63^, norepinephrine^64;65;66^, serotonin ^63^, and dopamine^67;68^ can also significantly effect CBF and the BOLD signal.

What do input parameters of an LTI model capture in BOLD fMRI? We show that an LTI system acts predominantly as a high-pass filter and highlights the rapid transient fluctuations in the BOLD signal. We provide evidence that the influence of sensory inputs is identifiable in the estimated inputs to sensory cortices. More importantly, the task-related changes that are temporally decoupled from the sensory stimuli, such as the motor cortex’s activation following the offset of visual cues and onset of behavioral outputs, are also captured as external inputs to the LTI system.

Prior research has reported brain-wide and heterogeneous task-related changes in the BOLD signal power spectrum^69;70;10^ and estimated system parameters^2;71^. However, we provide evidence that the time-varying unknown exogenous (i.e., extra-cortical) inputs also likely contribute to non-stationarities in the cortical dynamics. Specifically, we demonstrate in silico that determining the LTI system’s parameters from periods with unknown stimuli can lead to high estimation errors in system and input parameters. We verify these observations empirically by showing that LTI system parameters identified from resting-state, instead of task BOLD time series, result in notably more accurate identification of unknown extra-cortical inputs’ spatiotemporal profiles in task scans. Together, these results highlight the importance of modeling and interpreting the brain’s dynamic functional connectivity and non-stationarity as an open system

Can the brain during resting-state scans be fully described as a *linear* and *time-invariant* system? Prior studies demonstrate that temporal fluctuations in the BOLD signal (< 0.1 Hz) cannot be fully attributed to linear stochastic processes^72;73;74^, and suggest that the nonlinearities in the BOLD signal could be attributed to the presence of a strange attractor^73^. Additionally, other neuroimaging studies using paradigms such as “temporal summation” have more directly probed the *system* and provide evidence of system nonlinearities ^75;76;77;78^.

Model-based approaches such as work by^79;78^ have concluded that nonlinear transduction of rCBF to BOLD is sufficient to account for the nonlinear behaviors observed in the BOLD signal. However, care should be taken in the interpretation of these results as in the temporal summation framework, where the profile of input is assumed to be known and is approximated by an abstract stimulus representation. We believe our framework provides a novel avenue for testing the system linearities through the examination of the estimated unknown inputs in summation paradigms. Specifically, the delay between estimated and known external inputs can be further leveraged to tease out the nonlinear components of hemodynamic response function (e.g., vascular) from the neural impulse response function.

Stationary signals are characterized by time-invariant statistical properties, such as mean and variance^80^. To date, several tests have been proposed to examine the non-stationarity of BOLD time series and the presence of dynamic functional connectivity, including test statistics based on the variance of the FC time series^81;82^, the FC time series’ Fourier transform^83^, multivariate kurtosis of time series^27;28^, non-linear test statistics^47^, and wavelet-based methods^31;29^, among others^32^. These methods commonly compare measured properties between the time series of empirical data and a suitable surrogate or null time series that is designed to lack time-varying properties through non-parametric resampling^84;85^, phase-randomization^86;83^, or generative models^31;47^, and the choice of measured properties and null models profoundly impact on the outcomes of stationarity tests in conflicting reports on BOLD signal^29;30;27;28^.

Notably, the presence of non-stationarity in the outputs does not directly imply the underlying system’s non-stationarity. An LTI system’s outputs, for instance, while receiving non-stationary external inputs, can also display time-varying properties. As mentioned earlier, using internal system parameters of an LTI system estimated over resting-state scans enables more accurate identification of exogenous inputs’ spatiotemporal profile task scans. These results suggest that a large-scale stationarity model of the brain with time-varying external inputs can, in theory, account for a large portion of the observed task-related changes in cortical dynamics. It is worth noting that any possible task-related changes in the underlying system parameters are also captured as external inputs in an LTI framework. Therefore, from the system identification and model-fitting perspective, it is likely that a linear switching system with higher degrees of freedom would improve the fit. Beyond the goodness-of-fit of the model, care should be taken in interpreting the epiphenomenal large-scale models’ parameters and their changes at the micro-scale biophysical level.

However, the impetus for this work is to highlight the estimates’ notable sensitivity to the unknown, and thus, unaccounted external inputs. More practically, when simulated with a wideband unknown external inputs, our results suggest that an open LTI model estimated during restingstate allows us to uncover the influence of these unknown drivers of BOLD dynamics. Nevertheless, participants’ cortices receive external stimulation even during resting-state scans, contributing to estimation inaccuracy and the system’s outputs’ non-stationarity. In this work, we aim to disentangle the non-stationarity of the *system* from its *outputs* over resting-state by examining estimated inputs’ non-stationarity. Our results show that external inputs’ non-stationarity over resting-state scans are spatially inhomogeneous, with identified DMN regions showing the highest levels consistently across different analyses. These observations are in line with prior reports of higher dynamic functional connectivity of these brain structures over rest^47^. Despite the presence of possible confounding factors such as unaccounted nonlinearities and non-stationarities in the recording noise^87;88^, our framework and observations provide new insight into the external drivers of cortical dynamics and factors that contribute to their non-stationarity. Recent system-identification^89^ and control-theoretic^90^ work have also demonstrated the utility of a stationary system in explaining BOLD dynamics. Together these findings pave the way for principled model-based control of pathological brain dynamics, such as depression and schizophrenia, using open-loop external or closed-loop neurofeedback stimulation.

Historically, a narrow band of slow frequencies between 0.01 to 0.1 Hz was thought to contain information relevant to underlying neural activity, and that the higher frequency (> 0.1 Hz) BOLD activity considered mainly as an artifact^9;91^. Our results also demonstrate that the primary oscillatory modes of the LTI model of the resting-state BOLD display similar slow frequencies heterogeneously over the brain. In addition, the hemodynamic response function (HRF) is also expected to dampen the higher frequency neural activity significantly. More recent evidence, however, portrays a broadband picture of BOLD signal fluctuations with frequencies up to 0.25 Hz^92;93;10;13^ and even higher^14^. We also provide converging evidence that despite the expected low-pass filtering of HRF, information about the stimulus-related activity can still be extracted from the BOLD signal even as high as ≈ 0.4 Hz. Future work can leverage acquisition protocol with higher sampling rates than HCP and rapid stimuli capable of inducing brain-wide activations to accurately delineate the inputs’ attenuation profile by HRF at higher frequencies.

However, the HRF plays another critical role in biophysical models where it enables the approximation of the latent neural states from the BOLD signal. This is one of the main limitations of our simplified model, as it incorrectly assumes that the BOLD signal in one region (instead of the underlying neural activity) can cause changes in the BOLD signal in the connected regions. This assumption for spatially inhomogeneous HRF functions can, in theory, lead to incorrect identification of the external inputs’ focus and error in the direction and speed of the interactions within functional networks. We believe the overlapping patterns of inputs and the task activation maps identified using the conventional univariate general linear model analysis suggest that the above-mentioned error is likely tolerable. To improve the estimated unknown inputs’ accuracy, future work should leverage the formulated quantitative spatiotemporal^94;79;95;96^ models, or the more recent models informed by the precise mechanisms of neurovascular coupling^97^. Nevertheless, care should be taken in these or other related deconvolution-based inferences^98;99;100^, since as mentioned earlier, they rely on the assumption of a known profile of HRF or inputs. Future work can also leverage neural adaptation paradigms to influence the neural response timing and help tease out the neural and vascular components’ contributions to the modeled inputs.

Structured recording noise such as autocorrelated noise can negatively impact the modeled system^87;88^, and the estimated input. Although we have included global mean signal regression (GSR) ^42^ as a preprocessing step to account for the shared global noise that is present in many of the functional networks^101;102;103^, our model is unable to account for other unknown structured (e.g., autocorrelated) and time-varying recording noise^87;88^. Moreover, GSR may also introduce artifact, as in addition to the shared noise, it also removes any global activation patterns (e.g., vigilance^104^ or arousal^105^) and can alter the correlation structure. These limitations are the source of ongoing controversy around this noise reduction method ^106^. Having weighed the potential drawbacks of GSR against the major concerns regarding the significant global artifacts such as the cardiac and respiratory noise, we adopted this preprocessing step. Nevertheless, it would be beneficial to investigate the spectral profile of the global signal and the impact of GSR on the estimated system and inputs’ spectral characteristics.

One of the current limitations of our proposed framework is that the estimated inputs’ accuracy depends on the internal and recording noise levels. We show that group-level analysis and repeated measurement designs are effective strategies to increase signal-to-recording noise and to increase the estimated inputs’ accuracy. In addition, although we can not accurately tease out the contributions of internal noise from other sources of noise, our simulations and experimental results suggest lower levels of internal noise relative to external drivers in task fMRI. We draw this conclusion based on the relatively large input estimation errors associated with system parameters identified during external stimulation.

Finally, it is worth highlighting that model-based *data-driven* methods such as our proposed framework and the *hypothesis-driven* methods such as DCM^1^ are complementary approaches, suited for interrogation of different aspects of system and output dynamics. For instance, DCM can also be leveraged fruitfully for a more accurate estimation of the system, and consequently, external input parameters using highly controlled experimental designs with known external input profiles. Though, as mentioned before, care should be taken in the interpretation of the results produced by methods that incorporate priors, as the boxcar regressors commonly used to model the profile of external inputs are merely abstractions and do not account for other possible factors such as anticipatory responses, adaptation, or other unknown drivers that shape the profile of external inputs. However, data-driven approaches are particularly advantageous when the brain is driven by extensive complex inputs, for instance, during naturalistic stimuli (e.g., watching a movie), or in general, if we lack a priori information or hypothesis on the structure of external inputs – for instance, during the healthy resting-state or pathological brain activity such as epileptic discharges^107^.

## 5 Conclusions

We show that the proposed framework provides an avenue to uncover the structure of the unknown drivers of BOLD signal fluctuations and shines light on factors that contribute to its apparent non-stationarities. However, more significantly, our results highlight the importance of modeling and interpreting the brain’s dynamic functional connectivity as an *open* system. Broadly, our approach provides a framework for understanding the brain’s large-scale functional dynamics and non-stationarities, mechanistically via the modeled system and its time-varying drivers.

## 6 Data Availability

Data were provided [in part] by the Human Connectome Project, WU-Minn Consortium (Principal Investigators: David Van Essen and Kamil Ugurbil; 1U54MH091657) funded by the 16 NIH Institutes and Centers that support the NIH Blueprint for Neuroscience Research; and by the McDonnell Center for Systems Neuroscience at Washington University. All data analyzed in this manuscript came from part of the HCP S1200 release from the resources available in the public domain. All data used in this study are available for download from the Human Connectome Project (https://www.humanconnectome.org). The resting-state and task fMRI dataset are Open Access, therefore first-tier permission must be granted by the HCP to access the data. We provide list of all participants’ ID used in this study in supplementary table 1.

## 7 Code Availability

The costume scripts for estimating the LTI system and external input parameters are available at https://github.com/aashourv/LTI_BU.

## 8 Ethics Statement

All subject recruitment procedures and informed consents were approved by the Washington University Institutional Review Board (IRB). For more details see^35^.

## 9 Acknowledgements

A.A. was supported by The Mirowski Family Foundation. Inc. S.P. gratefully acknowledge the support by the National Science Foundation under the NSF award under Grant Number CMMI 1936578. D.S.B and B.L acknowledge support from the National Institute of Health (R01 NS099348-01) and The Neil and Barbara Smit Fund. D.S.B. also acknowledges support from the John D. and Catherine T. MacArthur Foundation, the Alfred P. Sloan Foundation, the ISI Foundation, the Paul Allen Foundation, the Army Research Laboratory (W911NF-10-2-0022), the Army Research Office (Bassett-W911NF-14-1-0679, Grafton-W911NF-16-1-0474, DCIST-W911NF-17-2-0181), the Office of Naval Research, the National Institute of Mental Health (2-R01-DC-009209-11, R01 - MH112847, R01-MH107235, R21-M MH-106799), the National Institute of Child Health and Human Development (1R01HD086888-01), and the National Science Foundation (BCS-1441502, BCS-1430087, NSF PHY-1554488 and BCS-1631550). Data were provided [in part] by the Human Connectome Project, WU-Minn Consortium (Principal Investigators: David Van Essen and Kamil Ugurbil; 1U54MH091657) funded by the 16 NIH Institutes and Centers that support the NIH Blueprint for Neuroscience Research; and by the McDonnell Center for Systems Neuroscience at Washington University.

## Author Contributions

A.A.: Conceptualization of this study, Methodology, Data Analysis, Writing. S.P.: Conceptualization of this study, Methodology, Writing. M.B.: Data Preprocessing, Writing. J.K.: Methodology, Writing. D.B.: Conceptualization of this study, Writing. B.L.: Conceptualization of this study, Writing.

## A Supplementary Information

In the Supplementary Information document, in addition to the supplementary figures, we provide details of our proposed joint-estimation algorithm (SI1),the LTI system’s output simulation (SI2), the dimensionality of the external inputs (SI3), determining the optimal number of eigenvector clusters (SI4), the sparsity of system matrix and external inputs (SI5), the effect of estimation window’s length on input’s accuracy (SI6), principal component analysis of LTI model’s residuals (SI7), and the list of de-identified HCP subjects used in this study in supplementary table 1.

